# Transcriptional regulation of sphingolipid metabolism in budding yeast

**DOI:** 10.1101/2021.11.05.467429

**Authors:** Nao Komatsu, Yuko Ishino, Rina Shirai, Ken-taro Sakata, Motohiro Tani, Tatsuya Maeda, Naotaka Tanaka, Mitsuaki Tabuchi

## Abstract

Global control for the synthesis of lipids constituting a bilayer of cell membranes is known to be with a small number of transcription factors called master transcriptional regulators, which target a wide range of genes encoding lipid metabolism enzymes and/or their regulators. Although master transcriptional regulators of glycerophospholipids and sterols have been identified in both yeast and mammals, this aspect of sphingolipid metabolism is not yet understood. In the present study, we identified the C2H2-type zinc finger transcription factor, Com2, as a master transcriptional regulator of sphingolipid metabolism in the budding yeast, *Saccharomyces cerevisiae*. The target of rapamycin complex 2 (TORC2)-activated protein kinase Ypk1 is known to regulate sphingolipid metabolism. Activated Ypk1 stimulates the activity of serine palmitoyl transferase (SPT), the first-step enzyme in sphingolipid biosynthesis, by phosphorylating and inhibiting Orm1/2, a negative regulator of SPT. This regulation of SPT activity is thought to be a major pathway in the regulation of sphingolipid metabolism. In the present study, we found that inhibition of sphingolipid synthesis upregulates the expression of Com2, which in turn leads to the concomitant expression of Ypk1. The upregulation of Ypk1 expression was found to be dependent on a putative Com2-binding site in the *YPK1* promoter. Our results also suggested that Com2 senses intracellular sphingolipid levels through a pathway independent of TORC2-Ypk1-mediated sensing of sphingolipids. Our results revealed an additional layer of mechanistic regulation that allows cells to maintain appropriate levels of sphingolipid biosynthesis and to rapidly induce this process in response to environmental stresses.

**Significance Statement:** One of the major regulatory mechanisms involved in the control of lipid metabolism in bilayers of biological membranes is regulation at the transcriptional level by master transcriptional regulators that control the transcription of genes encoding lipid metabolism enzymes and/or their regulators. In the present study, we identified the C2H2-type zinc finger transcription factor Com2 as a master transcriptional regulator in sphingolipid metabolism. We found that Com2 regulates sphingolipid metabolism by transcriptionally controlling the expression of Ypk1, which regulates Orm1/2, a negative regulator of serine palmitoyl transferase, the first-step enzyme in sphingolipid biosynthesis, through phosphorylation. Our study revealed a new layer of regulation that allows the maintenance of an appropriate level of sphingolipid biosynthesis for a rapid response to environmental stresses.

## Introduction

Cell membranes are composed of lipid bilayers, which consist of three major types of membrane lipids: glycerophospholipids, sphingolipids, and sterols (mainly cholesterol in mammals and ergosterol in yeasts) (1). The synthesis of lipid bilayers during membrane formation involves the coordinated assembly of multiple lipid species, which requires the coordination of different levels, including lipid synthesis, uptake, metabolism, and subcellular distribution (2). This homeostasis is important for the maintenance of cellular and tissue functions, and its dysfunction in humans can lead to numerous diseases, such as cancer or type-2 diabetes (3). The principal instruments for the global control of lipid metabolism are regulated by master transcriptional regulators, which target genes encoding metabolic enzymes and/or their regulators working in specific metabolic pathways (2).

For example, in yeast, the transcription factors Ino2/Ino4 function in the expression of several glycerophospholipid synthases (4). Under logarithmic growth or inositol-depleted conditions, the repressor Opi1 is bound to the endoplasmic reticulum (ER) membrane in a phosphatidic acid (PA)-dependent manner, and Ino2/Ino4 promote transcription. However, in steady-state or inositol-rich conditions, the decrease in PA on the ER membrane causes Opi1 to translocate to the nucleus, thereby inhibiting Ino2/Ino4-dependent transcription and repressing glycerophospholipid synthesis (5-9). In the cholesterol metabolism of animal cells, the membrane-bound transcription factors, SREBPs, are synthesized as inactive precursors that are anchored in the membrane of the ER under sterol-rich conditions (10). SREBPs are translocated to the Golgi apparatus in a COPII vesicle-dependent manner (11-13), where it is sequentially cleaved by Site-1 and Site-2 proteases, detached from the membrane (14-17), and translocated to the nucleus, thus promoting the expression of genes involved in cholesterol metabolism, such as HMG CoA reductase, the rate-limiting enzyme in cholesterol metabolism (2, 18). Thus, in glycerophospholipid and sterol metabolism, it is well known that the synthesis of these lipids is coordinately regulated by master transcriptional regulators, in both yeast and mammals. However, master transcriptional regulators of sphingolipid metabolism remain elusive (19). Sphingolipids and their metabolites play key cellular roles as both structural components of membranes and as signaling molecules that mediate responses to physiological cues (19). Recent studies using budding yeast, *Saccharomyces cerevisiae*, as a model system have improved our understanding of metabolic enzymes and how these enzymes are regulated to ensure sphingolipid homeostasis (19, 20). The regulation of sphingolipid metabolism by the target of rapamycin (TOR) complex 2 (TORC2)-Ypk1 pathway has been studied in detail in yeast (21-24). As a regulatory mechanism of sphingolipid biosynthesis, TORC2, in concert with Slm1/2, which was identified as a synthetic lethal gene along with *MSS4* encoding a phosphatidylinositol 4-phosphate 5-kinase (24, 25), phosphorylates Ypk1 upon depletion of complex sphingolipids at the plasma membrane (22, 23). Phosphorylated Ypk1 further phosphorylates Orm1/2, which negatively regulates serine palmitoyl transferase (SPT) (26-28), which is the first step in sphingolipid metabolism. Thereafter, the phosphorylation of Orm1/2 through Ypk1 results in an increase in SPT activity. The TORC2-Ypk1 signal has also been reported to activate ceramide synthase by phosphorylating the N-terminal sites of the ceramide synthase catalytic subunits, Lag1/Lac1 (29). Together, TORC2-Ypk1-dependent signaling simultaneously upregulates the two most important steps in the sphingolipid biosynthetic pathway.

To analyze the effect of the TORC2-Ypk1 pathway on reduced ceramide synthesis using the regulatory subunit mutant of ceramide synthase, *lip1-1*, we found that *lip1-1* is hypersensitive to myriocin (Myr), which specifically inhibits SPT (Fig. 1A) (30). To search for a new regulator of sphingolipid metabolism, we used a multi-copy suppressor of Myr sensitivity in *lip1-1* and identified Com2, a C2H2-type zinc finger transcription factor that has not been previously described in relation to sphingolipid metabolism. Further analysis revealed that the expression level of Com2 increased in response to a decrease in sphingolipid level upon Myr treatment and the expression level of Ypk1 and TORC2-mediated phosphorylation of Ypk1 increased synchronously with an increase in Com2 expression. In the Com2-deficient (*com2*Δ) strain, there was a decrease in both the Myr-dependent expression level of Ypk1 and its activation by TORC2, indicating that the upregulation of Ypk1 in response to decreased sphingolipid levels is Com2-dependent. In addition, a putative Com2-binding site (CBS) is present in the promoter region of *YPK1*, and the *P*_*YPK1*_*-CBS*Δ strain, in which this sequence was removed by means of genome editing, was Myr-sensitive, and canceled the Com2-dependent upregulation of Ypk1 upon Myr treatment. These results indicated that Com2 is a novel master transcriptional regulator that positively regulates sphingolipid metabolism by sensing a decrease in sphingolipid levels, increasing its expression at the transcriptional level, and promoting the expression of downstream *YPK1*, to stimulate sphingolipid synthesis.

**Figure 1.**
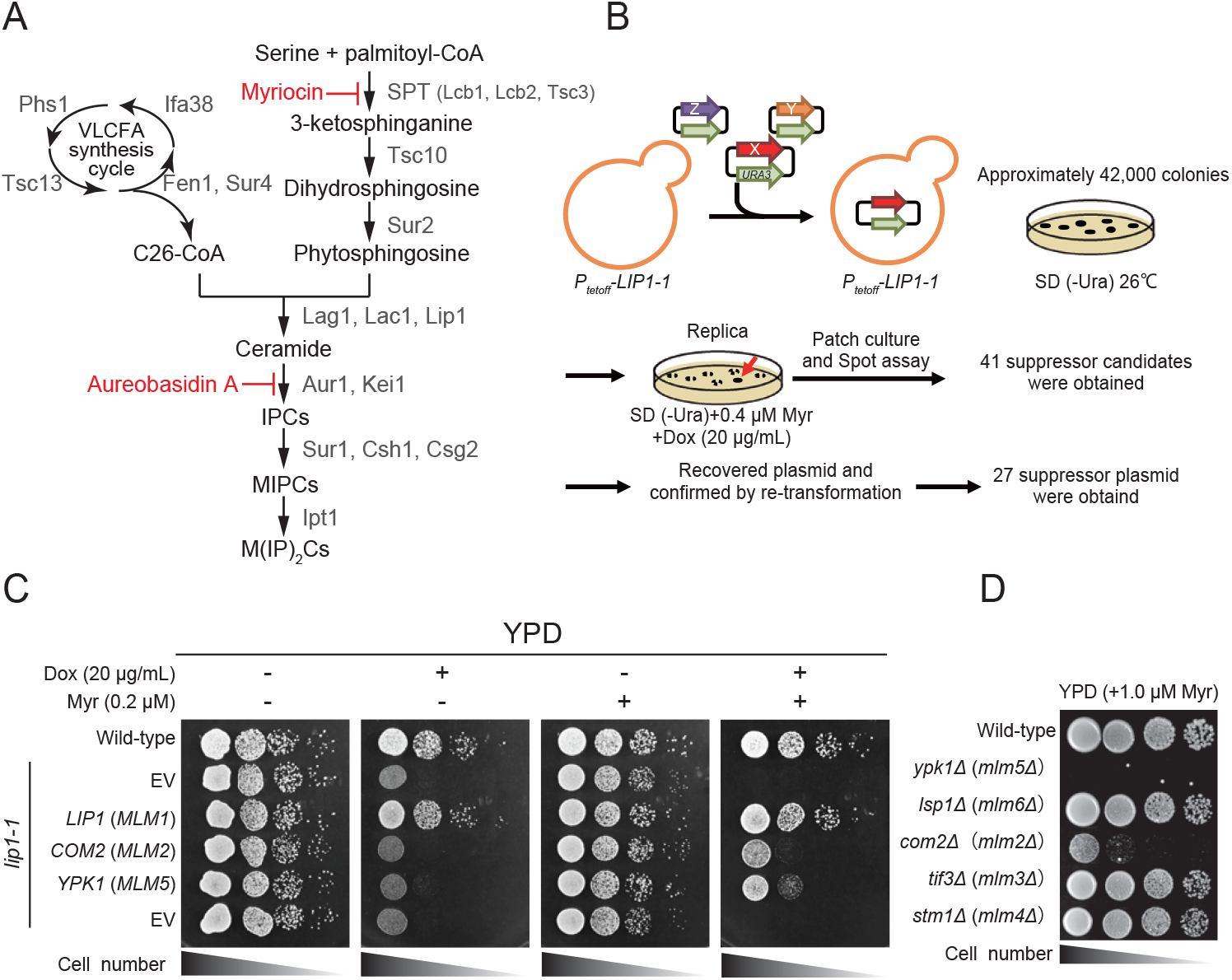
Screening for novel regulatory factors of sphingolipid metabolism. (*A*) *De novo* sphingolipid biosynthesis pathway in yeast, *Saccharomyces cerevisiae*. The pathway and proteins responsible for the synthesis of yeast sphingolipids are shown. IPC, inositol phosphorylceramide; MIPC, mannosylinositol phosphorylceramide; M(IP)_2_C, mannosyldiinositol phosphorylceramide. Myriocin (Myr) and aureobasidin A (AbA) inhibit the indicated steps of the sphingolipid biosynthesis pathway. (*B*) Schematic representation of screening for novel regulatory factors of sphingolipid metabolism by using the Myr-sensitive phenotype of *lip1-1* cells. (*C*) Wild-type and *lip1-1* cells carrying an indicated plasmid were spotted at a 10-fold serial dilution on YPD supplemented with (0.2 µM) or without Myr, in the presence (20 µg/mL) or absence of doxycycline (Dox). (*D*) The diploid homozygous knockout cells of *MLM* genes (*ypk1*Δ, *lsp1*Δ, *com2*Δ, *tif3*Δ, and *stm1*Δ) were spotted at a 10-fold serial dilution on YPD supplemented with 1.0 µM Myr.

## Results

### A screen for novel regulatory factors involved in sphingolipid metabolism

Previously, we generated and characterized a *lip1-1* strain in which ceramide synthesis decreased in the presence of doxycycline (Dox) (30). Dox-treated *lip1-1* cells are hypersensitive to Myr, an inhibitor of SPT, the first step in sphingolipid biosynthesis (Fig. 1A), presumably due to the synthetic effect of the Dox-dependent decrease in ceramide synthesis and inhibition of SPT. We suspected that *lip1-1* cells would become resistant to Myr if a protein that facilitates sphingolipid biosynthesis was overexpressed in these cells. Therefore, to identify novel regulatory factors for sphingolipid metabolism, *lip1-1* cells were transformed with a multi-copy genomic library, following which the transformants that grew on the media with Myr were selected (Fig. 1B). From the ∼42,000 transformants that were screened, we identified 27 plasmids that allowed *lip1-1* cells to grow on Myr-containing media (Figs. 1B and S1). These plasmids were sequenced and classified into 12 groups (12 *MLM* genes: <u>M</u>ulti-copy suppressor for *<u>L</u>ip1-1* <u>M</u>yr-sensitive phenotype, Table S1) according to the genes they contained. Among these, the *YPK1* gene (*MLM5*), which encodes for a protein kinase downstream of the TORC2 signaling pathway, which is known to be involved in the regulation of sphingolipid metabolism (23, 29), was identified, thus validating that genes involved in the regulation of sphingolipid metabolism can be obtained by means of this screening. Besides the *LIP1* gene (*MLM1*), the *COM2* gene (*MLM2*), which encodes a C2H2-type zinc finger transcription factor, was the most abundant (Fig. 1C, Table S1).

To clarify whether the obtained *MLM* genes are involved in the regulation of sphingolipid metabolism, we examined the Myr sensitivity of each gene-knockout homozygous diploid strain. Upon doing so, we found that *com2*Δ, as well as, *ypk1*Δ exhibited Myr sensitivity, while the other *MLM* gene-deficient strains did not (Figs. 1D and S2).

These results suggested that Com2 might play a role as a regulatory factor in sphingolipid metabolism, and we further investigated this in the next part of the study.

### Com2 functions upstream of the TORC2-Ypk1 pathway

Overexpression of Com2 or Ypk1 suppressed the Myr sensitivity of *lip1-1* (Fig. 1C) and deletion of any of these genes conferred Myr sensitivity (Fig. 1D), suggesting that these two proteins function in the same pathway involved in the regulation of sphingolipid metabolism. Com2 is a C2H2-type zinc finger transcription factor consisting of 443 amino acid residues with two zinc fingers at its C-terminus. In addition to that, several putative phosphorylation sites for AGC kinases, including Ypk1, were found in its amino acid sequence (Figs. 2A and S4). Based on these observations, we hypothesized that Com2 is involved in sphingolipid metabolism downstream of Ypk1 and that Ypk1 impacts the phosphorylation and function of Com2.

**Figure 2.**
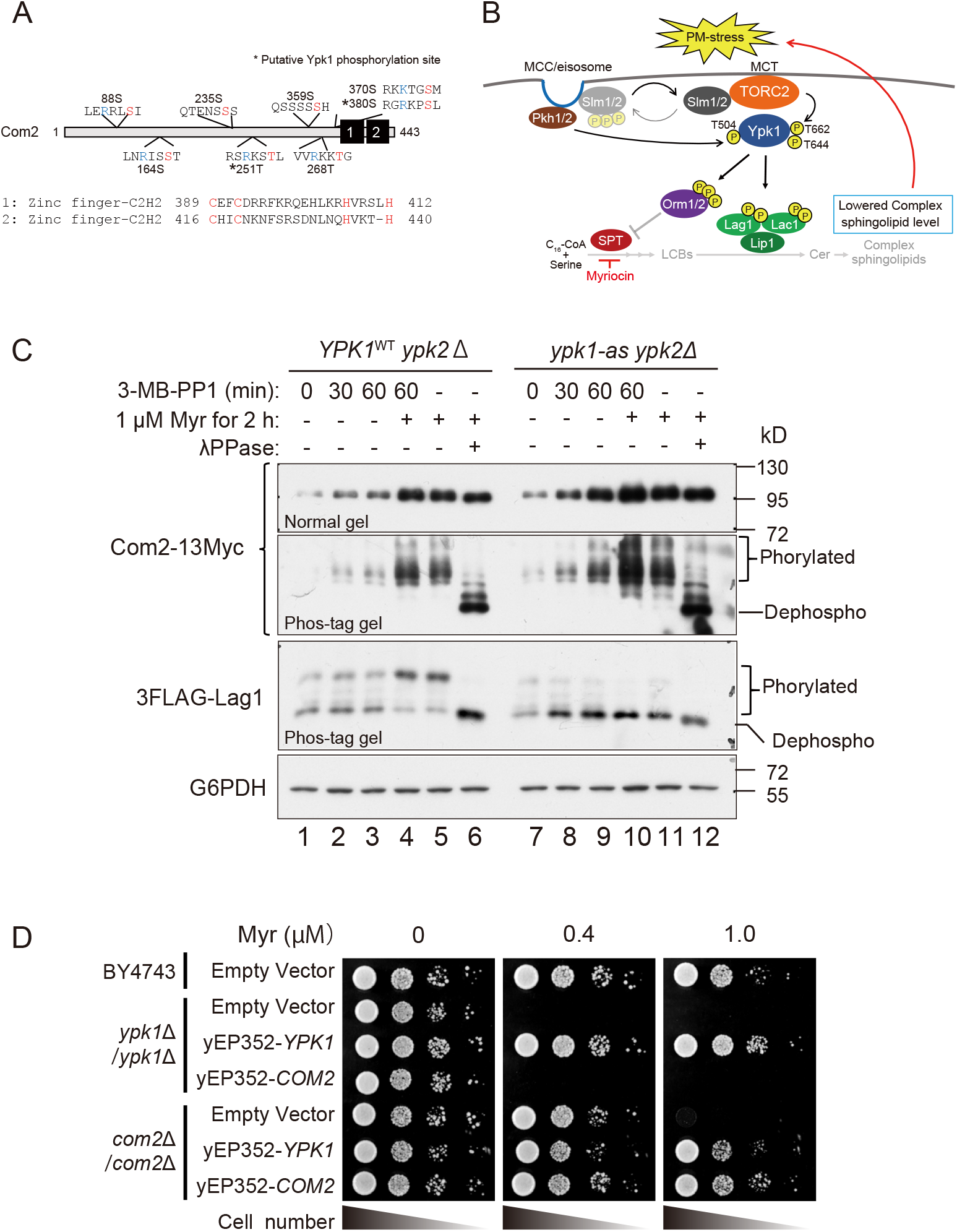
Com2 functions upstream of the TORC2-Ypk1 pathway. (*A*) Schematic representation of Com2, with the positions of the predicted AGC-kinase-dependent phosphorylation sites and two Zinc-finger domains highlighted. (*B*) Schematic representation of the TORC2-Ypk1-dependent regulatory mechanism of sphingolipid biosynthesis. (*C*) Wild-type *YPK1*^WT^ *ypk2*Δ cells and analogue-sensitive mutant *YPK1*^L424A^*ypk2*Δ cells carrying the plasmid expressing Com2-13Myc were treated with 3MB-PP1 for the indicated time-periods and then treated with 1.0 µM Myr for 2 h. The lysates were then resolved using phos-tag or normal SDS-PAGE and immunoblotted with anti-Myc, anti-FLAG, or anti-G6PDH antibodies, to detect Com2-13Myc, 3FLAG-Lag1, or G6PDH (loading control), respectively. (*D*) Wild-type (BY4743) and knockout homozygote cells of *ypk1*Δ or *com2*Δ carrying an indicated multi-copy plasmid were spotted at a 10-fold serial dilution on YPD supplemented with the indicated concentrations of Myr.

To test this hypothesis, we investigated the effect of limiting Ypk1 activity (Fig. 2B) on the phosphorylation state of Com2. To this end, we employed *ypk1-as ypk2*Δ cells, which express an ATP analog-sensitive (AS) allele, Ypk1(L424A), and titrated down its activity by addition of an efficacious inhibitor, 1-(tert-butyl)-3-(3-methylbenzyl)-1H-pyrazole[3,4-d]pyrimidin-4-amoine (3-MB-PP1), which has no effect on the wild-type cells (Fig. S3A) (22, 27, 29). As previously reported (29, 30), Myr treatment stimulated the phosphorylation of 3FLAG-tagged Lag1 in the wild-type cells, as verified using phos-tag SDS-PAGE (Fig. 2C; lanes 1-5 in the 3FLAG-Lag1 blot), but this effect was deficient in *ypk1-as ypk2*Δ cells (Fig. 2C; lanes 7-11 in the 3FLAG-Lag1 blot), confirming the inhibition of Ypk1 activity. It is noted that the *ypk1-as ypk2*Δ mutant did not show Myr-dependent phosphorylation of Lag1 even in the absence of 3-MB-PP1 and was also sensitive to Myr (Fig. 3B), suggesting that this mutant appears to be a hypomorphic allele and treatment of 3-MB-PP1 completely block Ypk1 activity. In contrast to phosphorylation of Lag1, a 13Myc epitope-tagged Com2 was highly phosphorylated in both the wild-type and *ypk1-as ypk2*Δ cells (Fig. 2C), indicating that the phosphorylation state of Com2 is not dependent on Ypk1 activity. Interestingly, Com2 expression levels were increased upon Myr treatment in the wild-type cells (Fig. 2C; lanes 1-3 *vs*. lanes 4-6 in the Com2-13Myc blot), and this effect was more prominent in the 3-MB-PP1-treated *ypk1-as ypk2*Δ cells (Fig. 2C; lanes 7-9 *vs*. lanes 10-12 in the Com2-13Myc blot).

**Figure 3.**
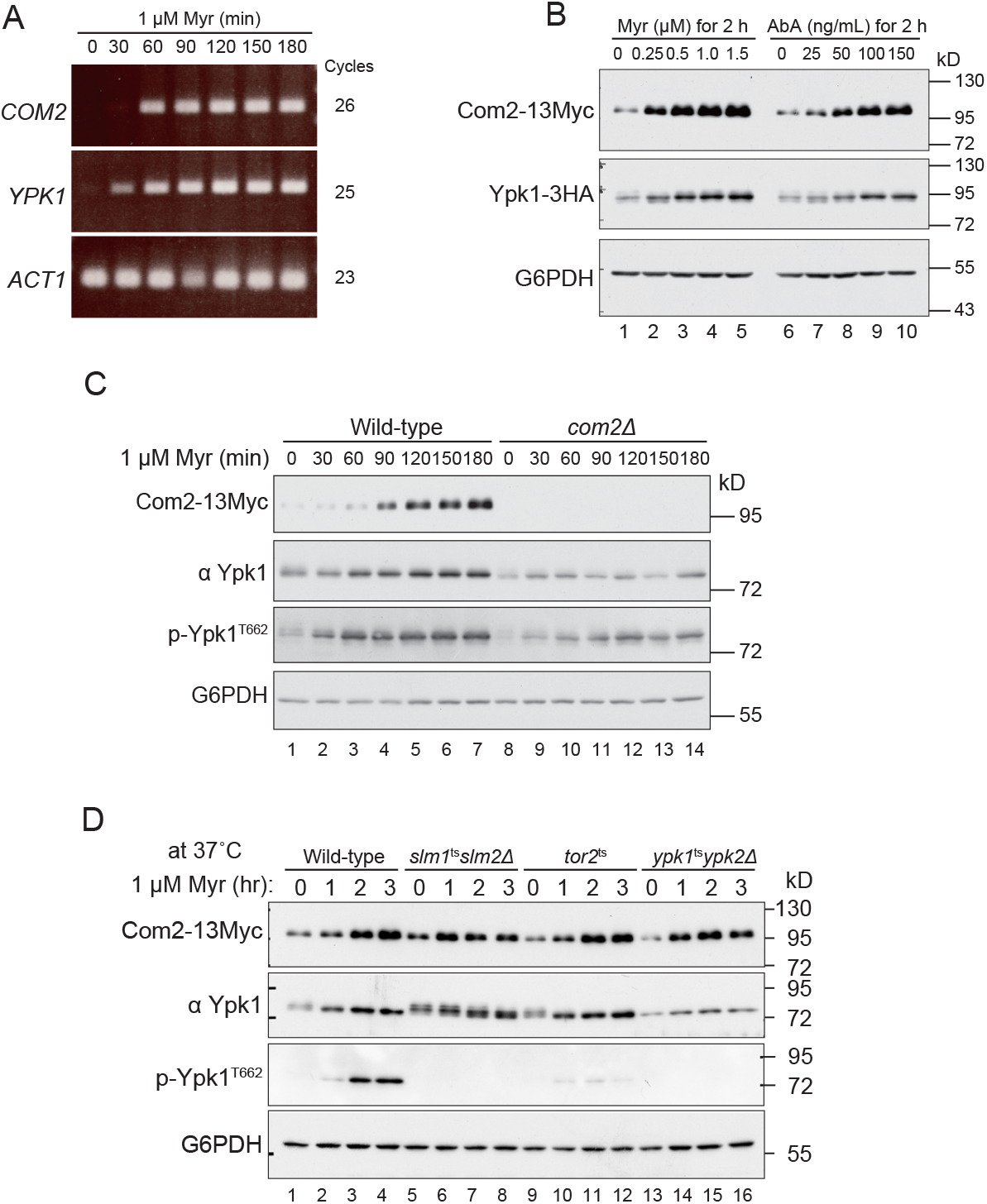
Ypk1 is expressed in a Com2-dependent manner. (*A*) RT-PCR detection of *COM2* and *YPK1* mRNA. Wild-type cells were grown to mid-log phase in SD liquid medium and treated with 1.0 µM Myr. Total RNA was prepared from the cells harvested at the indicated time-points and subjected to RT-PCR using *COM2-, YPK1-*, and *ACT1*-specific primers. (*B*) Wild-type (*COM2*-13Myc *YPK1*-3HA) cells were grown to mid-log phase in SD liquid medium and treated with or without the indicated concentrations of Myr or AbA for 2 h. The cells were then harvested and total cell lysates were prepared from them. The lysates were resolved using SDS-PAGE and immunoblotted with anti-Myc, anti-HA, or anti-G6PDH antibodies, to detect Com2-13Myc, Ypk1-3HA, or G6PDH (loading control), respectively. (*C*) Wild-type and *com2*Δ cells were grown to mid-log phase in SD liquid medium and then treated with 1.0 µM Myr. The cells were then harvested at the indicated time-point and total cell lysates were prepared from them. The lysates resolved using SDS-PAGE and immunoblotted with anti-Ypk1, anti-Myc, anti-G6PDH antibodies, or phospho-specific antibodies, to detect Ypk1, Com2-13Myc, G6PDH (loading control), or a TORC2-dependent phosphorylation site (hydrophobic motif) at the 662^th^ threonine (T662) of Ypk1/2, respectively. (*D*) Wild-type, *slm1*^ts^*slm2*Δ, *tor2*^ts^, and *ypk1*^ts^*ypk2*Δ cells were grown to mid-log phase in SD liquid medium at 26°C and then shifted to 37°C for 30 min and treated with 1.0 μM Myr. The cells were then harvested at the indicated time-point and total cell lysates were prepared from them. The lysates were resolved using SDS-PAGE and immunoblotted with anti-Ypk1, anti-Myc, anti-G6PDH antibodies, or anti-phospho-specific antibodies, to detect Ypk1, Com2-13Myc, G6PDH (loading control), or TORC2-dependent phosphorylation of Ypk1/2, respectively.

Next, we performed an epistatic analysis between *COM2* and *YPK1* genes, for which we confirmed the suppression of Myr sensitivity in each gene-deletion strain by overexpressing each gene using multi-copy plasmids. Overexpression of *COM2* did not suppress the Myr sensitivity of *ypk1*Δ cells, whereas overexpression of *YPK1* suppressed the Myr sensitivity of *com2*Δ cells (Fig. 2D), indicating that Com2 functions upstream of Ypk1.

Taken together, these results suggested that Com2 functions upstream of the TORC2-Ypk1 pathway.

### Com2 expression increases in response to a decrease in sphingolipids

We observed that Myr treatment increased the expression of Com2 in both the wild-type and *ypk1-as ypk2*Δ cells (Fig. 2C), which prompted us to investigate the effect of Com2 expression in the inhibition of sphingolipid biosynthesis in more detail. In addition, since Com2 has been suggested to function upstream of Ypk1, the expression of Ypk1 was also investigated. Wild-type cells were treated with Myr for different time-points, and *COM2* and *YPK1* mRNAs were detected in the total RNA preparations using reverse transcription-PCR (RT-PCR). Surprisingly, Myr treatment increased the transcriptional levels of both *COM2* and *YPK1* mRNAs in a time-dependent manner (Fig. 3A). Furthermore, western blot analysis revealed that the expression levels of both Com2 and Ypk1 were dose-dependently upregulated upon treatment with not only Myr, but also aureobasidin A (AbA) (Fig. 3B), which is a potent inhibitor of inositol phosphoceramide synthase (Fig. 1A), indicating that these increases were caused by a decrease in the levels of complex sphingolipids. Since Com2 is a transcription factor that functions upstream of Ypk1, we examined whether Com2 directly affects the expression of Ypk1. We performed a time-course experiment to determine the expression levels of Com2 and Ypk1 during Myr treatment in wild-type and *com2*Δ cells and confirmed the results by means of western blot. There was a time-dependent increase in the expression level of Com2 upon Myr treatment in wild-type cells, accompanied by increases in not only the expression of Ypk1 but also TORC2-dependent phosphorylation of the hydrophobic motif of Ypk1 (p-Ypk1 T662) (Fig. 3C; lanes 1-7). On the other hand, in *com2*Δ cells, there was a decrease in both the upregulation of Ypk1 and Myr-dependent phosphorylation of Ypk1 T662 observed in wild-type cells (Fig. 3C; lanes 1-7 *vs*. lanes 8-14), indicating that these increases are dependent on Myr-induced expression of Com2.

TORC2 senses the plasma membrane stress associated with a decrease in the level of complex sphingolipids in the plasma membrane *via* eisosome-associated adaptor proteins, Slm1/2, and then activates Ypk1 to stimulate sphingolipid synthesis (Fig. 2B). Next, we investigated the contribution of the Slm-TORC2-Ypk1 pathway to the Myr-induced expression of Com2. We observed Myr-induced expression of Com2 even after blocking of the Slm-TORC2-Ypk1 pathway at non-permissive temperatures using each temperature-sensitive mutant (Fig. 3D), indicating that the induction of Com2 expression upon decreased complex sphingolipids is due to a sensing pathway that is independent from the Slm-TORC2-Ypk1 pathway.

Taken together, we demonstrated that Com2 is upregulated in response to a decrease in complex sphingolipids, accompanied by upregulation of Ypk1 expression.

### Overexpression of Com2 causes increased expression of Ypk1 *via* a putative CBS in the *YPK1* promoter

To confirm in detail that Com2 functions upstream of Ypk1, the promoter region of *COM2* on the chromosome was replaced with a tet-off promoter by means of homologous recombination, to generate a *P*_*tet-off*_*-GFP-COM2* strain, in which Com2 expression can be regulated by Dox (Fig. 4A). Using this strain, we first examined the Myr sensitivity of yeast cells when GFP-Com2 was overexpressed or repressed. The yeast cells overexpressing GFP-Com2 (Dox^-^) exhibited resistance to Myr, as compared to the wild-type cells (Fig. 4B, upper panel), while the yeast cells with repressed expression of GFP-Com2 (Dox^+^) were as sensitive to Myr as *com2*Δ cells (Fig. 4B, lower panel).

**Figure 4.**
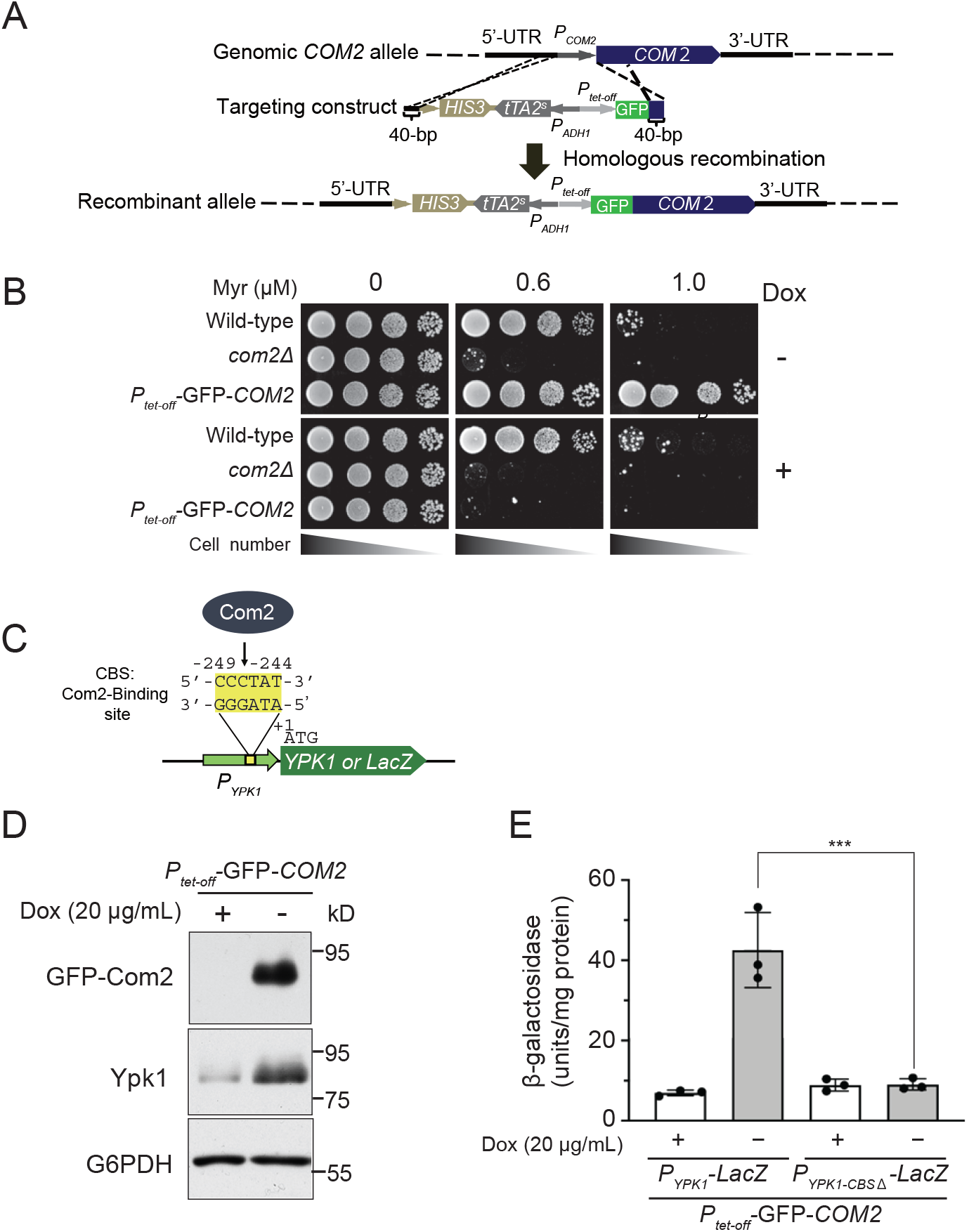
Overexpression of Com2 causes increased expression of Ypk1. (*A*) The chromosomal *COM2* promoter (*P*_*COM2*_) was replaced by a tet-off promoter and GFP was fused to the N-terminus of Com2, to generate the *P*_*tet-off*_-GFP-*COM2* strain. (*B*) Wild-type, *com2*Δ, and *P*_*tet-off*_-GFP-*COM2* cells were spotted at a 10-fold serial dilution on YPD supplemented with 0.6 or 1.0 µM Myr, in the presence (20 µg/mL) (+) or absence (-) of Dox. (*C*) *YPK1* possesses a putative Com2-binding site (CBS) in the promoter region. (*D*) *P*_*tet-off*_-GFP-*COM2* cells were grown to mid-log phase in SD liquid medium, in the presence (+) or absence (-) of Dox, and then treated with or without 1.0 µM Myr for 2 h. The lysates were resolved using SDS-PAGE and immunoblotted with anti-Com2, anti-Ypk1, anti-Myc, or anti-G6PDH antibodies, to detect GFP-Com2, Ypk1, or G6PDH (loading control), respectively. (*E*) *P*_*tet-off*_-GFP-*COM2* cells carrying p*P*_*YPK1*_*-LacZ* or p*P*_*YPK1-CBSΔ*_*-LacZ* plasmid were grown to mid-log phase in SD liquid medium, in the presence (+) or absence (-) of Dox. Lysates were obtained from the indicated samples and assayed for β-galactosidase activity. Values have been represented as mean±SD (n≥3). \*\*\**P*<0.001, as assessed using one-way ANOVA, with Tukey–Kramer multiple comparison tests.

Because the promoter region of *YPK1* contains a putative Com2-binding site (CBS), 5’-CCCTAT-3’ (Fig. 4C) (31), we investigated whether Com2 is directly involved in the regulation of the expression of *YPK1*. To this end, we examined the effect of Com2 overexpression on the expression levels of Ypk1. As shown in Fig. 4D, the expression level of Ypk1 was markedly increased under GFP-Com2-overexpressed condition (Dox^-^). Furthermore, we constructed a *lacZ* reporter containing the -493-bp region of the *YPK1* promoter sequence (Fig. 4C) and measured *YPK1* promoter activity using β-galactosidase activity as an indicator. We observed an approximately six-fold increase in transcription under the GFP-Com2-overexpressed condition (Dox^-^), as compared to that under the GFP-Com2-repressed condition (Dox^+^) (Fig. 4E). Importantly, this effect was eliminated by deleting the CBS sequence from the *YPK1* promoter (Fig. 4E), indicating that Com2 directly stimulates the expression of *YPK1 via* the CBS sequence.

Taken together, these results demonstrated that Com2 is involved in sphingolipid metabolism, presumably through regulation of Ypk1 transcription and expression.

### Com2 promotes sphingolipid synthesis, partly through the expression of Ypk1

To investigate whether Com2 is directly involved in the transcription and expression of *YPK1*, we generated *P*_*YPK1*_-*Com2-binding site*Δ (*P*_*YPK1*_*-CBS*Δ) mutant cells that lack the putative CBS in the promoter region of chromosomal *YPK1*, by means of CRISPR-Cas9-mediated genome editing (Fig. 5A).

**Figure 5.**
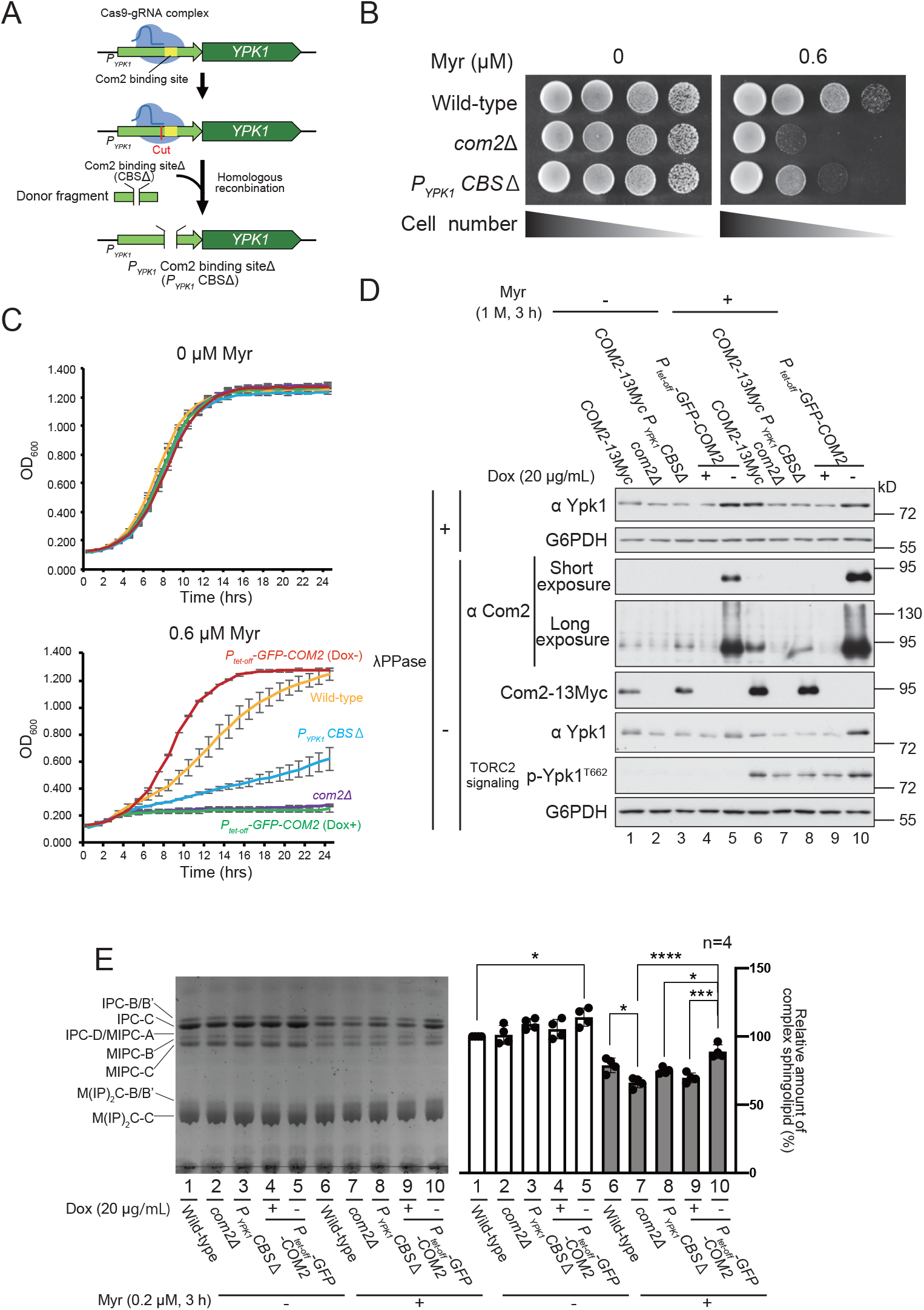
The Com2-binding site in the *YPK1* promoter is required for both proper expression and TORC2-dependent phosphorylation of Ypk1. (*A*) A schematic representation of the generation method of the *P*_*YPK1*_-*Com2 binding site*-deleted (*P*_*YPK1*_-*CBS*Δ) strain using CRISPR-Cas9-mediated genome editing. (*B*) Wild-type, *com2*Δ, and *P*_*YPK1*_*-CBS*Δ cells were spotted at a 10-fold serial dilution on YPD supplemented with or without 0.6 µM Myr. (C) Wild-type, *com2*Δ, *P*_*YPK1*_-*CBS*Δ, and *P*_*tet-off*_-GFP-*COM2* cells were grown to the exponential phase, diluted to OD_600_=0.1, and grown in YPD liquid medium treated with or without 0.6 µM Myr, in the absence (-) or presence (+) of Dox (20 µg/mL), in a microtiter plate. (*D*) *COM2*-13Myc, *com2*Δ, *P*_*YPK1*_*-CBS*Δ *COM2*-13Myc, and *P*_*tet-off*_-GFP-*COM2* cells were grown to mid-log phase in SD liquid medium, in the presence (+) or absence (-) of Dox (20 µg/mL) and treated with 1.0 µM Myr for 3 h. The lysates were resolved by means of SDS-PAGE and immunoblotted with anti-Ypk1, anti-Com2, anti-Myc, anti-G6PDH antibodies, or phosphospecific antibodies, to detect Ypk1, GFP-Com2, Com2-13Myc, G6PDH (loading control), or the hydrophobic motif (T662) of Ypk1/2, respectively. (*E*) TLC analysis of sphingolipids extracted from wild-type, *com2*Δ, *P*_*YPK1*_-*CBS*Δ, and *P*_*tet-off*_-GFP-*COM2* cells. Cells were grown to mid-log phase in YPD liquid medium, in the presence (+) or absence (-) of Dox (20 µg/mL), and treated with or without 0.2 µM Myr for 3 h. Complex sphingolipids were analyzed using TLC (left). The level of total complex sphingolipids in the wild-type cells was taken as 100% and each value is displayed as a graph (right). The data have been represented as mean±SD of four independent experiments. Asterisks denote statistically significant differences based on one-way ANOVA followed by Dunnett’s test (\**p*<0.05, \*\*\**p*=0.0002, and \*\*\*\**p*<0.0001).

First, we examined the Myr sensitivity of *P*_*YPK1*_-*CBS*Δ cells. The *P*_*YPK1*_*-CBS*Δ cells exhibited sensitivity to Myr, as compared to wild-type cells, but were not as sensitive as *com*2Δ or Dox-treated *P*_*tet-off*_*-GFP-COM2* (Dox^+^) cells (Fig. 5B-C).

Next, we examined the expression level of Ypk1 in *P*_*YPK1*_-*CBS*Δ cells using western blot. Since Ypk1 is detected as a smeared band on the electrophoresis gels because its N-terminus is also phosphorylated by Fpk1/Fpk2 (32), we dephosphorylated the sample by treating it with λPPase, to detect Ypk1 and compare the density of the bands more clearly. While the expression of Com2-13Myc in wild-type and *P*_*YPK1*_-*CBS*Δ cells was almost identical under Myr-treated conditions, the expression of Ypk1 was reduced in *P*_*YPK1*_-*CBS*Δ cells, as compared to that in wild-type cells (Fig. 5D; lane 6 *vs*. lane 8) and was comparable to that in *com2*Δ or Dox-treated *P*_*tet-off*_*-GFP-COM2* cells (Fig. 5D; lane 8 *vs*. lane 7 or 9). Similarly, there was a decrease in TORC2-dependent phosphorylation of the hydrophobic motif (p-Ypk1 T662) in *P*_*YPK1*_-*CBS*Δ cells, as compared to that in wild-type cells (Fig. 5D; lane 6 *vs*. lane 8), but comparable to that in *com2*Δ or Dox-treated *P*_*tet-off*_*-GFP-COM2* cells (Fig. 5D; lane 8 *vs*. lane 7 or 9).

Next, we performed TLC analysis to determine whether changes in Com2 expression affected the synthesis of sphingolipids. Consistent with the Myr-resistant phenotype, overexpression of GFP-Com2 (Dox^-^) resulted in a significant increase in sphingolipid levels under both normal (Fig. 5E; lane 1 *vs*. lane 5) and Myr-treated conditions (Fig. 5E; lane 6 *vs*. lane 10). On the other hand, upon Myr treatment, a significant decrease in sphingolipid levels was observed in *com2*Δ or GFP-Com2-repressed (Dox^+^) cells, as compared to that in wild-type cells (Fig. 5E; lane 6 *vs*. lane 7 or 9). The level of sphingolipids in Myr-treated *P*_*YPK1*_-*CBS*Δ cells was significantly lower, as compared to that in GFP-Com2-overexpressing (Dox^-^) cells (Fig. 5E; lane 8 *vs*. lane 10). However, we observed only a slight reduction in sphingolipid levels in Myr-treated *P*_*YPK1*_*-CBS*Δ cells, as compared to that in wild-type cells (Fig. 5E; lane 6 *vs*. lane 8), which is consistent with the fact that *P*_*YPK1*_-*CBS*Δ cells were not as sensitive to Myr as *com2*Δ cells (Fig. 5B). These results suggested that Ypk1 is a major target of Com2 under sphingolipid-depleted conditions, but other targets that regulate cellular sphingolipid levels must also exist.

In summary, our data demonstrated that Com2 increased the expression of Ypk1 through the CBS in the *YPK1* promoter, which partly contributes to the promotion of sphingolipid synthesis.

## Discussion

Bilayer synthesis during membrane formation requires the coordinated combination of multiple lipid species to regulate the levels of lipid synthesis, uptake, metabolism, and subcellular distribution (2). One of the major regulatory mechanisms for global control of lipid metabolism employs master transcriptional regulators that control the transcription of genes encoding lipid metabolic enzymes and their regulators. Master transcriptional regulators have been identified for glycerophospholipids and sterols in both yeast and mammals (2). However, our understanding of the transcriptional control of sphingolipid metabolism remains poor (19).

In this study, we demonstrated that Com2 is a master transcriptional regulator of sphingolipid metabolism by sensing the decrease in sphingolipid levels, upregulating its expression at the transcriptional level, and promoting Ypk1 expression to stimulate sphingolipid synthesis in budding yeast (Fig. 6).

**Figure 6.**
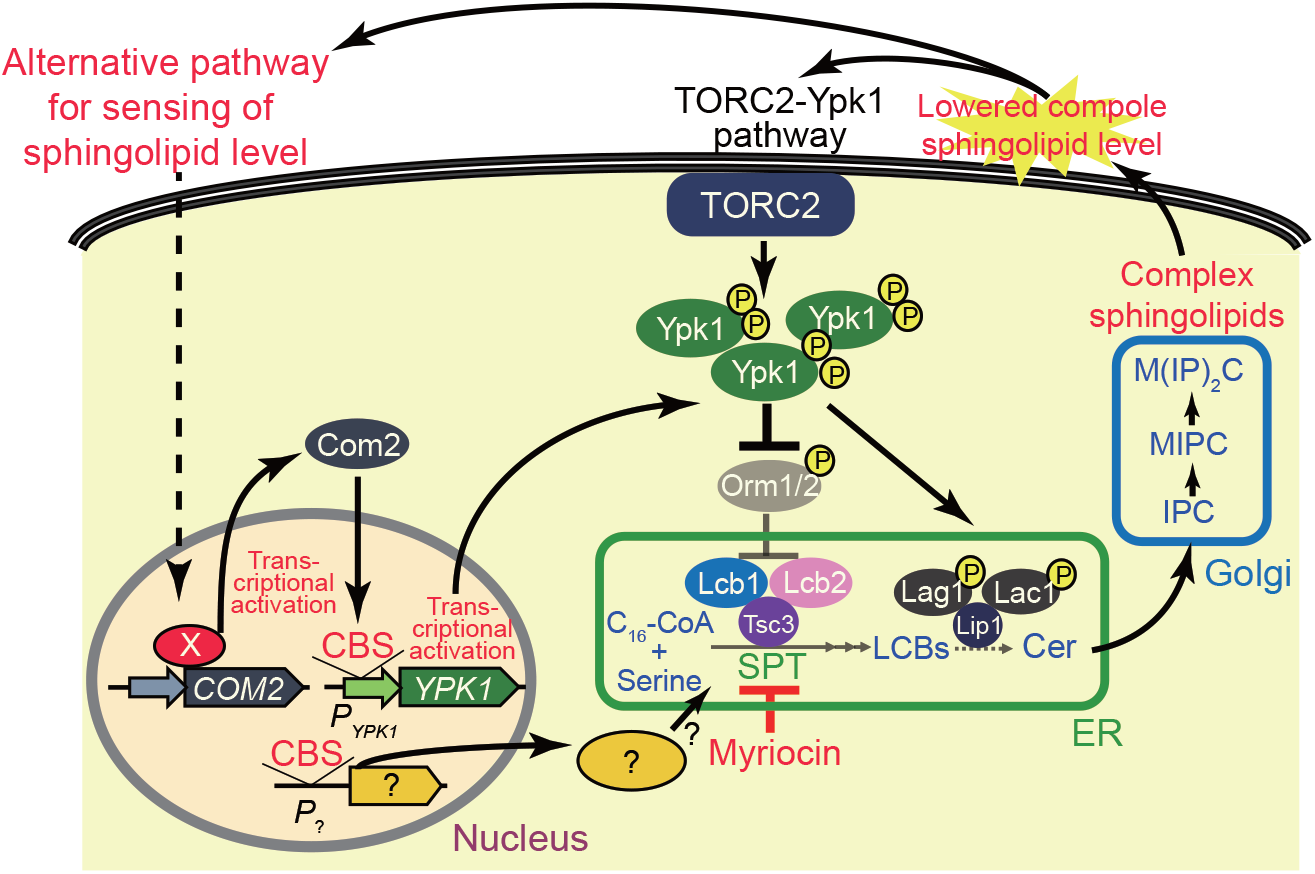
A model depicting the regulatory mechanism of sphingolipid biosynthesis *via* Com2-dependent transcriptional regulation. See more detail in Discussion.

Since there are at least eight putative phosphorylation sites for AGC kinase in the primary structure of Com2, as assessed using Computer search (Figs. 2A and S4), Com2 was initially presumed to function downstream of Ypk1. However, limiting Ypk1 activity did not affect the phosphorylation state of Com2 (Fig. 2C). Furthermore, in the mutant in which serine and threonine residues were replaced with alanine residues at all eight putative phosphorylation sites of Com2, there was a decrease in the phosphorylation of these mutants, but no significant decrease or increase in Com2 function, as evaluated by complementation of the Myr-sensitive phenotype of *com2*Δ cells (Fig. S4). The significance of Com2 phosphorylation needs to be clarified by identifying the critical phosphorylation sites and kinases that are responsible for phosphorylation and function of Com2.

Lage *et al*. (33) showed that *com2*Δ cells are sensitive to sulfur dioxide (SO_2_) and that Com2 is a major transcription factor that regulates the expression of genes involved in SO_2_ tolerance. In addition, Com2 has been identified as a transcription factor that controls the activity of UASru, a specific region within the *IME1* promoter that, when deleted, significantly reduces the transcription of *IME1*, the master regulator of meiosis in budding yeast (34). The relationship between Com2-dependent regulation of these pathways and enhancement of sphingolipid synthesis revealed in this study is unclear, but there may be a common mechanism in these different response pathways.

In the present study, we found that the major target of Com2 in sphingolipid metabolism was Ypk1, and the expression level of Ypk1 in *P*_*YPK1*_*-CBS*Δ cells was reduced to the same level as that in *com2*Δ or GFP-Com2-repressed cells (Fig. 5D), but these mutant cells were not as sensitive to Myr (Fig. 5B). In addition, overexpression of GFP-Com2 in *P*_*YPK1*_*-CBS*Δ cells conferred resistance to Myr and enhanced sphingolipid synthesis without increasing Ypk1 expression and TORC2-dependent phosphorylation of Ypk1 (Fig. S5A-C). These results suggested that Com2 has other targets that regulate sphingolipid metabolism, in addition to Ypk1. We searched for potential Com2 target genes in the *S. cerevisiae* genome based on the CBS consensus motif (31) and found the CBS sequence in the promoter of *LCB1* (Fig S6), which encodes for one of the catalytic subunits of SPT (35). Interestingly, the *lacZ* reporter fused with the *LCB1* promoter revealed that the transcription of this promoter increased approximately 2.7-fold upon Com2 overexpression, and this increase was canceled by the deletion of the CBS sequence (Fig. S6C), suggesting that *LCB1* is one of the targets of Com2. In addition to *LCB1*, several other candidate genes were found to be associated with sphingolipid metabolism (data not shown), and further analysis of these genes is expected to reveal the full extent of Com2-mediated regulation of sphingolipid metabolism at the transcriptional level.

A key issue of future research is to understand how cells modulate the expression of Com2 in response to sphingolipid levels. In particular, what is the physical basis for sphingolipid sensingã Com2 expression was increased at the mRNA level when complex sphingolipids were reduced upon Myr treatment. Intriguingly, the induction of Com2 expression by inhibition of sphingolipid synthesis was also observed when the *COM2* promoter was replaced with the tet-off promoter (Fig. 5D; lane 5 *vs*. lane 10 and Fig. S5A; lanes 4, 5 *vs*. lanes 9, 10), suggesting that the induction of Com2 expression by inhibition of sphingolipid synthesis is promoter-independent. TORC2 senses the changes in membrane tension associated with a decrease in the level of complex sphingolipids in the plasma membrane *via* eisosome-associated adaptor proteins, Slm proteins, and then activates Ypk1 to stimulate sphingolipid synthesis *via* inhibition of Orm proteins and activation of ceramide synthases (Fig. 6) (22, 23, 29). Since Myr-induced Com2 expression was not affected by blocking the Slm-TORC2-Ypk pathway (Fig. 3D), it suggests that induction of Com2 expression upon decreased sphingolipid synthesis is due to an independent alternative pathway that is distinct from the TORC2-mediated sensing of sphingolipids (Fig. 6). Future studies are expected to elucidate the physical molecular basis of sphingolipid sensing, especially by elucidating the molecular mechanisms involved in the induction of Com2 expression.

## Materials and Methods

### Strains, plasmids, and media

Descriptions of the strains and plasmids used in this study are presented in Tables S2 and S3. *Escherichia coli* DH5α or JM109 was used as the bacterial host for plasmid construction. Coding sequences were amplified by means of PCR, using KOD -Plus-Neo polymerase (Toyobo) or PrimeSTAR^®^ GXL polymerase (TaKaRa Bio Inc.). The plasmids were sequenced to ensure that no mutations were introduced owing to the manipulation. Mutant constructs were generated using site-directed mutagenesis and confirmed using sequencing. The media used for yeast culture were YPD (1% yeast extract, 2% peptone, and 2% glucose) or synthetic dextrose (SD) medium (2% glucose and 0.67% yeast nitrogen base without amino acids). Appropriate amino acids and bases were added to the SD medium, as necessary. Yeast cells were cultured at 26°C, unless otherwise stated. All deletion and tagged strains were constructed using homologous recombination with PCR-generated DNA fragments, as described previously (36).

### Yeast growth assays

Plate growth and liquid growth assays were performed using a microplate reader, as previously described (30).

### Antibody production

The rabbit polyclonal antibody that recognizes Com2 was produced as follows. A His_6_-fused Com2 protein was expressed in *E. coli* Rosetta-gami B (DE3) pLysS cells and the total cell lysate was separated on an SDS-PAGE gel. The band corresponding to the His_6_-Com2 fusion protein was excised from the SDS-PAGE gel and electroeluted from the gel slices using an AE-3590 electrochamber (Atto). An aliquot of approximately 300 μg of the fusion protein was emulsified with Freund’s complete adjuvant and injected intramuscularly or subcutaneously into young female Japanese white rabbits. Antisera were purified using an affinity column immobilized with His_6_-ProS2-Com2 (382-443 a.a.) fusion protein.

### Western blot

Protein extracts were prepared using the TCA precipitation method (30), loaded onto normal or phos-tag SDS-PAGE gels, and transferred to PVDF membranes (Fujifilm Wako). λPPase treatment was performed as described previously (30). The primary antibodies and dilutions used in this study were 1:5,000 αCom2, 1:10,000 α-Myc (MBL, My3, M192-3), 1:10,000 α-HA (Roche, 3F10, 11867423001), 1:5,000 α-FLAG (Sigma-Aldrich, M2, F1804), 1:1,000 αYpk1, 1:5,000 αphospho-Ypk2 (T659) (30) for phospho-Ypk1 (T662) and phospho-Ypk2 (T659), and 1:50,000 αG6PDH (Sigma-Aldrich, A9521).

### Sphingolipid analysis

Sphingolipids were extracted from *S. cerevisiae* and analyzed using TLC, as described previously (37).

### Total RNA isolation and RT-PCR analysis

Logarithmic-phase wild-type cells grown in SD liquid medium were treated with 1.0 μM Myr for the indicated time and harvested. Following that, total RNA was isolated from the cells using Sepasol (Nacalai Tesque). cDNA was synthesized from 1.0 μg of total RNA using Rever Tra Ace™ (Toyobo) primed with Oligo (dT) 20 primer. The primers used for RT-PCR were 5’-CAACGCCAATGCCAAGGAAA-3’ and 5’-ATCCTGTTCAGCCCGTCAAG-3’ for *COM2*; 5’-GTTCCACCGCCACATAAGGA-3’ and 5’-GCTTCTCGTCTTGCAAACCC-3’ for *YPK1*; and 5’-GCCTTCTACGTTTCCATCCA-3’ and 5’-GGCCAAATCGATTCTCAAAA-3’ for *ACT1*.

### Construction of *LacZ* reporter plasmids

The fragments containing the promoter regions of *YPK1* and *LCB1* were amplified by means of PCR and cloned into the XhoI and BamHI sites of the 4×*CDRE-LacZ* fusion plasmid, pAMS366 (38) (gifted by Dr. Martha Cyert, Stanford University) using the In-Fusion^®^ Cloning Kit (Clontech), resulting in *P*_*YPK1*_*-LacZ* and *P*_*LCB1*_*-LacZ* reporter plasmids, respectively.

### Genome editing for generation of the *P*_*YPK1*_*-CBSΔ* strain

The pMT1603 plasmid, constructed from the pSpCas9(BB)-2A-GFP plasmid (39) combined with the Z_4_EV expression system (40), co-expresses the nuclease protein Cas9 and gRNA, which guides the Cas9 protein with the target promoter sequence of *YPK1* (Table S3). The repair DNA fragment was amplified from pNK19 carrying the *P*_*YPK1*_*-CBSΔ* fragment (Table S3). The yeast wild-type strain was co-transformed with the pMT1603 plasmid and the repair DNA fragment using the lithium acetate transformation method. After selecting the uracil prototrophic transformants, the genomic DNA purified from the yeast transformants was used for genotyping by means of PCR using specific primers, to confirm the deletion of the CBS in the chromosomal *YPK1* promoter. The pMT1603 plasmid was lost by means of 5-FOA counter selection.

### Statistical analysis

Statistical analysis was performed using Prism 8 (GraphPad) or Excel (Microsoft) software, and the specific tests carried out have been mentioned in the text and figure legends.

## Acknowledgments

We express our deep appreciation to Prof. Scott D. Emr for generously providing the plasmids and strains. We thank Profs. Isamu Kameshita and Noriyuki Sueyoshi for providing the λPPase-expressing plasmid. We also thank Prof. Martha Cyert for providing the *LacZ* reporter plasmid, pAMS366. We also appreciate Prof. Takayuki Sekito for his technical advice on the experimental methods. We acknowledge the technical expertise of the DNA Core Facility of the Gene Research Center, Kagawa University. This work was supported by JSPS KAKENHI grant numbers JP19K05828 (to M.Tabuchi), 20H03251 (to T.M.), 21H02118 (to M.Tani), and grants from the Ohsumi Frontier Science Foundation (to M.Tani and T.M.) and HUSM Grant-in-Aid (to T.M.).

## Notes

### Competing Interest Statement

The authors have declared no competing interest.

