## Supplemental Figures and Tables for "Transcriptional regulation of sphingolipid metabolism in budding yeast"

### Supplementary Information for
### Transcriptional regulation of sphingolipid metabolism in budding yeast

Nao Komatsu[1*], Yuko Ishino[1*], Rina Shirai[1], Ken-taro Sakata[1], Motohiro Tani[2], Tatsuya Maeda[3], Naotaka Tanaka[1] and Mitsuaki Tabuchi[1, 4]

Mitsuaki Tabuchi


**This PDF file includes:**

Figures S1 to S6
Tables S1 to S3
SI References

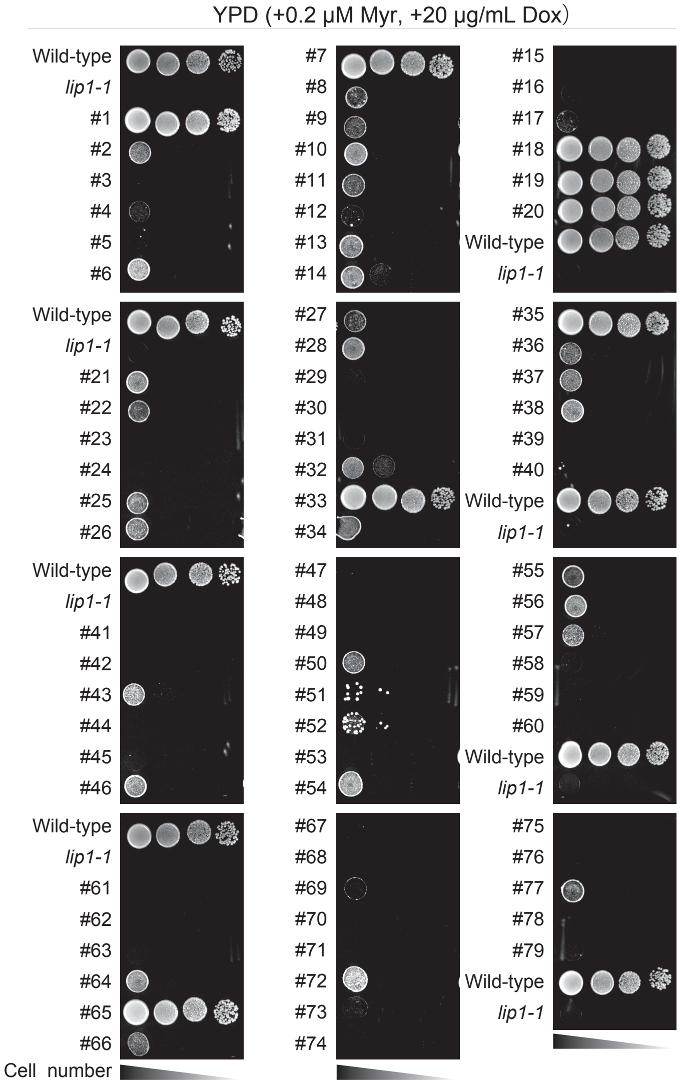


*MLM1* (*LIP1*) : #1, #7, #18, #19, #20, #33, #35, #65
*MLM2* (*COM2*) : #21, #34, #38, #46, #55, #64, #77
*MLM3* (*TIF3*) : #11, #13
*MLM4* (*STM1*) : #50, #54
*MLM5* (*YPK1*) : #6
*MLM6* (*LSP1*) : #9
*MLM7* (*BMH2*) : #17
*MLM8* (*RIM20* (partial), *CAF20*, *HEM4*, *RFM1*) : #26
*MLM9* (*TVP18*): #28
*MLM10* (*SRO77* (partial), *PKC1* (partial)): #36
*MLM11* (*PIN4*): #56
*MLM12* (*SUT2*): #66

**Fig. S1.** A screening for novel regulatory factors involved in sphingolipid metabolism using *lip1-1* mutant. Wild-type cells and *lip1-1* cells carrying each suppresser plasmid were spotted in 10-fold serial dilution on YPD supplemented with 0.2 μM Myriocin (Myr) in the presence of doxycycline (Dox) (20 μg/mL) and incubated for 3 d at 26℃.

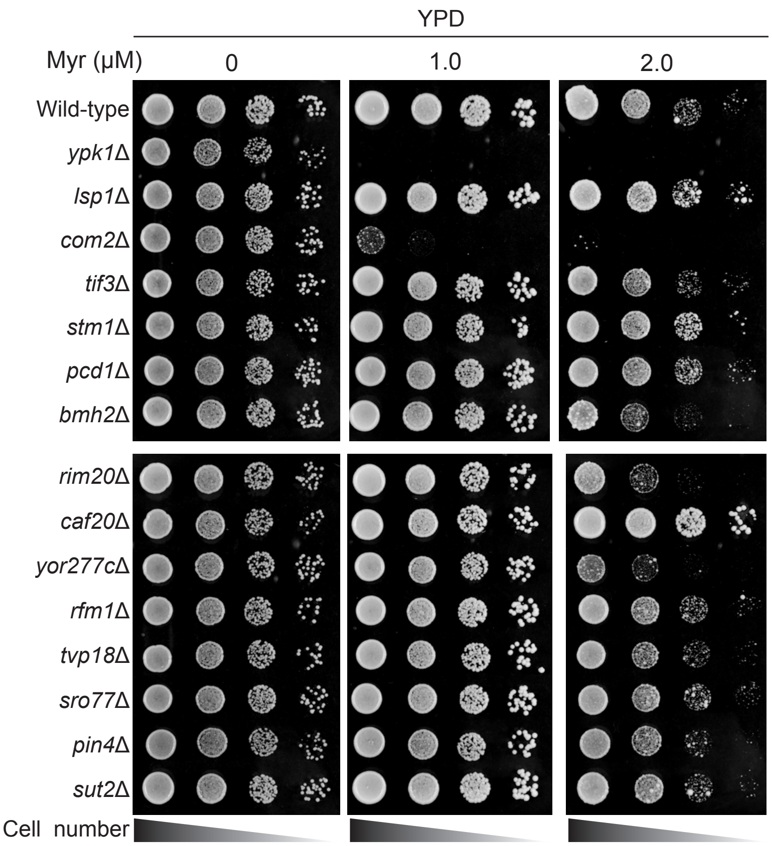


**Fig. S2.** Confirmation of the Myr sensitivity of the gene knockout homozygous diploid strain of obtained suppressor genes. Each gene knockout homozygous diploid cells of *MLM* suppressor genes were spotted in 10-fold serial dilution on YPD supplemented with indicated concentrations of Myr and incubated for 3 d at 26℃.

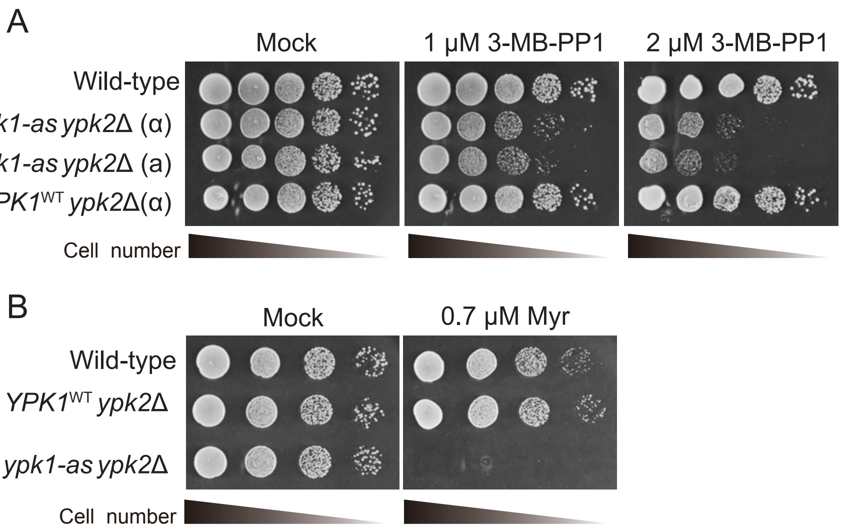


**Fig. S3.** Characterization of *ypk1-as ypk2*Δ mutant. (*A*) Wild-type, the ATP-analogue sensitive allele *ypk1*$^{L424A}$, *ypk1-as ypk2*Δ cells and the wild-type allele of Ypk1, *YPK1*$^{WT}$ *ypk2*Δ cells were spotted on YPD containing DMSO (Mock) or indicated concentration of 3MB-PP1 and incubated for 3 d at 26℃. *ypk1-as ypk2*Δ cells exhibit growth defect by limiting of Ypk1-activity by addition of 3MB-PP1. (B) Wild-type cells, *YPK1*$^{WT}$ *ypk2Δ* cells and *ypk1-as ypk2*Δ cells are spotted on YPD and YPD containing 0.7 μM myriocin and incubated for 2 d at 26℃. *ypk1-as ypk2*Δ cells exhibit Myr-sensitive without inhibition by addition of 3MB-PP1.

A

| Site | Sequence | Predicted phosphorylation sites: SGK1 | SGK3 | SGK | AGC | Phospho-site mutants: ST-A3 | ST-A5 | ST-A6 | ST-A7 | ST-A8 |
|---|---|---|---|---|---|---|---|---|---|---|
| 88S | NQLERRLSISDYRTE | | + | | + | | + | + | + | + |
| 164S | DGLNRISSTNNLKNL | | + | | | | + | + | + | + |
| 235S | PIQTENSSSQKMFKN | | | + | | | | + | + | + |
| 251T | FFRSRKSTLIKSLPL | + | + | + | + | + | + | + | + | + |
| 359S | SVQSSSSSHGLVVRK | | | + | | | | | + | + |
| 368T | GLVVRKKTGSMQKTR | | | | + | | | | | + |
| 370S | VVRKKTGSMQKTRGR | + | + | + | + | + | + | + | + | + |
| 380S | KTRGRKPSLIPDASK | + | | + | + | + | + | + | + | + |

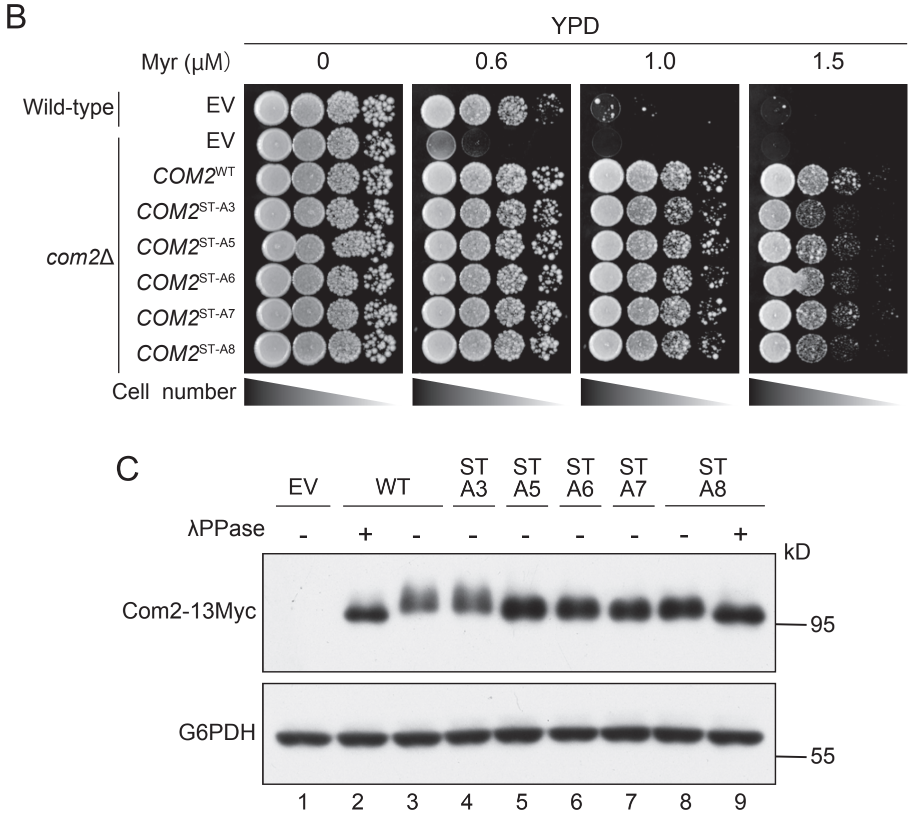


**Fig. S4.** Analysis of the putative phosphorylation sites of Com2. (*A*) The putative phosphorylation sites of Com2 by AGC kinases and the representation of mutation sites of phospho-site mutants. The phosphorylation sites were predicted by GPS web server: http://gps.biocuckoo.org/online.php . (*B*) Wild-type and *com2Δ* cells carrying an indicated plasmid were spotted in 10-fold serial dilution on YPD supplemented with indicated concentrations of Myr and incubated for 3 d at 26℃. (*C*) Wild-type carrying empty vector (EV) and an indicated plasmid cells were grown to mid-log in YNB (-Ura) liquid medium. Cells were then harvested and total cell lysates were prepared. The lysates resolved by SDS-PAGE and immunoblotted with anti-Myc or -G6PDH antibodies to detect Com2-13Myc or G6PDH to detect Com2-13Myc or G6PDH (loading control).

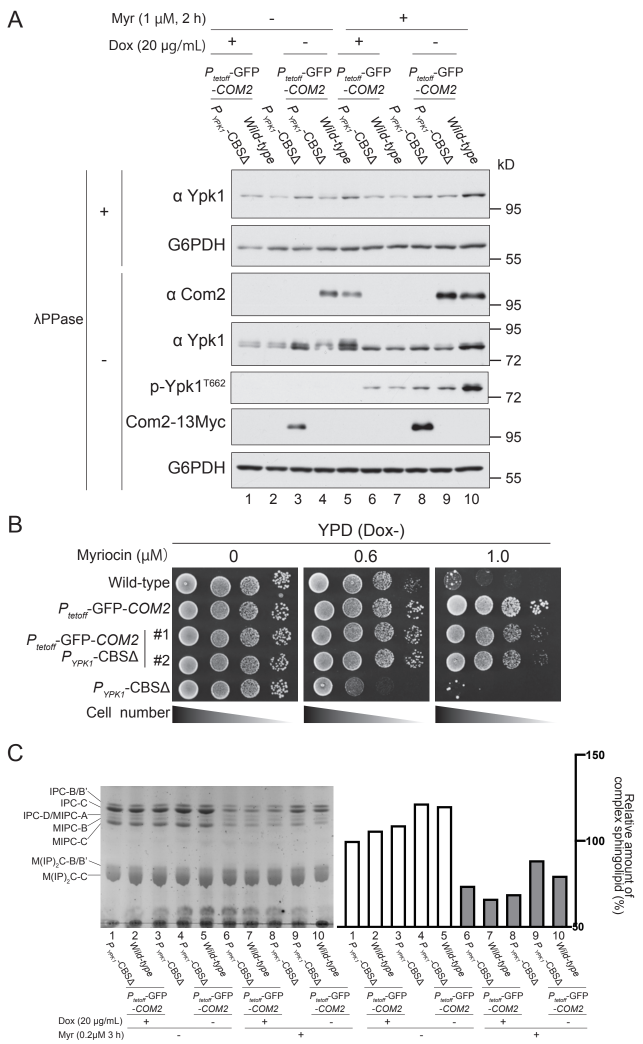


**Fig. S5.** Overexpression of GFP-Com2 confers Myr-resistant to $P_{YPK1}$-*CBS*Δ cells. (*A*) $P_{tet\text{-}off}$-GFP-*COM2* $P_{YPK1}$-*CBS*Δ, $P_{tet\text{-}off}$-GFP-*COM2* and $P_{YPK1}$-*CBS*Δ *COM2*-13Myc cells were grown to mid-log in SD liquid medium in the presence (20 μg/mL) (+) or absence (-) of Dox and treated with or without 1 μM Myr for 2 h. The lysates resolved by SDS-PAGE and immunoblotted with anti-Com2, -Ypk1, -Myc, -G6PDH or phospho-specific antibodies, to detect GFP-Com2, Ypk1, Com2-13Myc, G6PDH (loading control) or hydrophobic motif (T662) of Ypk1/2, respectively. (*B*) Wild-type, $P_{tet\text{-}off}$-GFP-*COM2*, $P_{tet\text{-}off}$-GFP-*COM2* $P_{YPK1}$-CBSΔ and $P_{YPK1}$-CBSΔ cells were spotted in 10-fold serial dilution on YPD supplemented with 0.6 or 1.0 μM Myr and incubated for 3 d at 26℃. (*C*) TLC analysis of sphingolipids extracted from $P_{tet\text{-}off}$-GFP-*COM2* $P_{YPK1}$-*CBS*Δ, $P_{tet\text{-}off}$-GFP-*COM2* and $P_{YPK1}$-*CBS*Δ cells. Cells were grown to mid-log in YPD liquid medium in the presence (+) or absence (-) of Dox (20 μg/mL) and treated with or without 0.2 μM Myr for 3 h. Complex sphingolipids were analyzed by TLC. The level of complex sphingolipids (IPCs, MIPCs and $M(IP)_2Cs$) in wild-type was taken as 100% and each value of indicated strains is displayed as a graph.

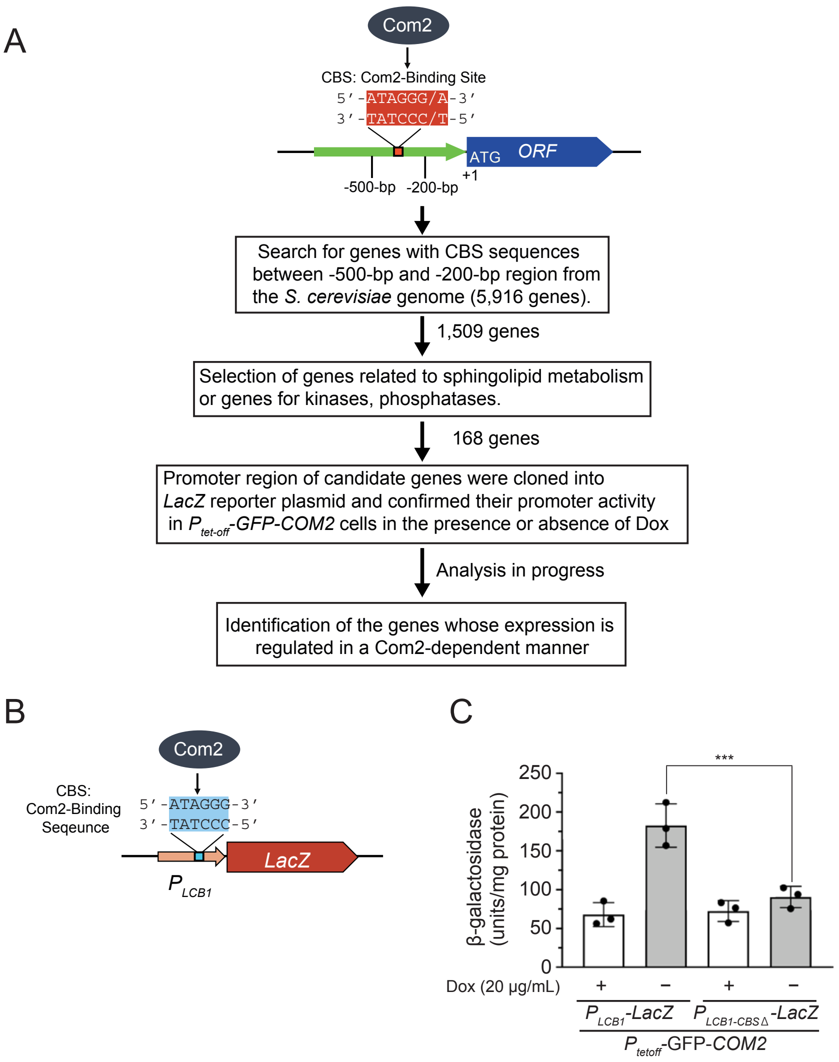


**Fig. S6.** *In silico* analysis to identify the candidate genes of the Com2 dependent expression in response to reduced sphingolipids. (*A*) Schematic representation of the workflow for the bioinformatic approach towards identifying Com2 targets from the *S. cerevisiae* genome using Yeast Genome Pattern Matching (https://www.yeastgenome.org/nph-patmatch). (*B*) *LCB1* promoter that possess a putative Com2 binding site (CBS) was fused to *LacZ* reporter. (*C*) $P_{tet\text{-}off}$-GFP-*COM2* cells carrying p$P_{YPK1}$-*LacZ* or p$P_{YPK1\text{-}CBS\Delta}$-*LacZ* plasmid were grown to mid-log in SD liquid medium in the presence (+) or absence (-) of Dox. Lysates were obtained from indicated samples and assayed for β-galactosidase activity. Values represent the mean ± SD (n≥3). ***$P$<0.001, one-way ANOVA post-test (Turkey-Kramer multiple comparison tests).

**Table S1.**

| *MLM* number | Number of Mutants | Genes | Functions |
|---|---|---|---|
| *MLM1* | 8 | *LIP1* | Ceramide synthase subunit |
| *MLM2* | 7 | *COM2* | Transcriptional factor that binds *IME1* upstream activation signal (UAS)ru |
| *MLM3* | 2 | *TIF3* | Transcription initiation factor eIF-4B |
| *MLM4* | 2 | *YLR149C-A* | Dubious open reading frame |
| | | *STM1* | Protein required for optimal translation under nutrient stress |
| *MLM5* | 1 | *YPK1* | S/T protein kinase |
| *MLM6* | 1 | *LSP1* | Eisosome core component |
| *MLM7* | 1 | *BMH2* | 14-3-3 protein |
| *MLM8* | 1 | *RIM20* | Protein involved in proteolytic activation of Rim101p |
| | | *CAF20* | Phosphoprotein of the mRNA cap-binding complex |
| | | *YOR277C* | Dubious open reading frame; almost completely overlaps the verified gene CAF20 |
| | | *SNR31* | H/ACA box small nucleolar RNA (snoRNA) |
| | | *SNR5* | H/ACA box small nucleolar RNA (snoRNA) |
| | | *HEM4* | Uroporphyrinogen Ⅲ synthase |
| | | *RFM1* | Component of the Sum1p-Rfm1p-Hst1p complex |
| *MLM9* | 1 | *MOT3* | Transcriptional repressor, activator |
| | | *TVP18* | Integral membrane protein |
| | | *ABF2* | Mitochondrial DNA-binding protein |
| *MLM10* | 1 | *SRO77* | Protein with roles in exocytosis and cation homeostasis |
| | | *PKC1* | Protein serine/threonine kinase |
| *MLM11* | 1 | *PIN4* | Protein involved in G2/M phase progression and response to DNA damage |
| | | *SEC17* | Alpha-SNAP cochaperone |
| *MLM12* | 1 | *SUT2* | Zn2Cys6 family transcriptional factor |

**Table S2. Yeast strains used in this study**

| Strain | Genotype | Reference or source |
|---|---|---|
| SEY6210 | *MATα leu2-3, 112 ura3-52 his3-Δ200 trp1-Δ901 lys2-801 suc2-Δ9* | (1) |
| SEY6210.1 | *MATa leu2-3, 112 ura3-52 his3-Δ200 trp1-Δ901 lys2-801 suc2-Δ9* | (1) |
| BY4741 | *MATa his3Δ1 leu2Δ0 met15Δ0 ura3Δ0* | Research Genetics |
| BY4742 | *MATα his3Δ1 leu2Δ0 lys2Δ0 ura3Δ0* | Research Genetics |
| BY4743 | *MATα/a* BY4741/BY4742 | Research Genetics |
| MTY1000 | SET6210; *HIS3MX6*::$P_{tetoff}$-*GFP-COM2* | This study |
| MTY1009 | SEY6210; *COM2-13MYC::HIS3MX6* | This study |
| MTY1025 | SEY6210; *ypk1Δ*::*HIS3MX6 ypk2Δ*::*HIS3MX6* pRS415-*YPK1*$^{WT}$ *COM2-13MYC::HIS3MX6* | This study |
| MTY1049 | SEY6210; *ypk1Δ*::*HIS3MX6 ypk2Δ*::*HIS3MX6* pRS415-*YPK1*$^{L424A}$ *COM2-13MYC::HIS3MX6* | This study |
| MTY1050 | SEY6210.1; *ypk1Δ*::*HIS3MX6 ypk2Δ*::*HIS3MX6* pRS415-*YPK1*$^{L424A}$ *COM2-13MYC::HIS3MX6* | This study |
| MTY1087 | SEY6210; $P_{YPK1}$*CBSΔ* | This study |
| MTY1192 | SEY6210; *tor2Δ*::*HIS3* YCplac111-*tor-21*$^{ts}$ *COM2-13MYC::HIS3MX6* | (2), This study |
| MTY1194 | SEY6210; *ypk1*$^{ts}$::*HIS3 ypk2Δ*::*HIS3MX6 COM2-13MYC::HIS3MX6* | (3) , This study |
| MTY1198 | SEY6210; *slm1Δ*::*HIS3MX6 slm2Δ*::*HIS3MX6* pRS415-*slm1-1* *COM2-13MYC::HIS3MX6* | (4) , This study |
| DYY1 | SEY6210; *HIS3MX6*::$P_{tetoff}$-*lip1-1* | (5) |
| YIY36 | SEY6210; *com2Δ*::*KANMX4* | This study |
| YIY50 | SEY6210; *COM2-13MYC*::*HIS3MX6 YPK1-3HA::KANMX4* | This study |
| NKY101 | SEY6210; *HIS3MX6*::$P_{tetoff}$-*GFP-COM2* $P_{YPK1}$*CBSΔ* | This study |
| NKY103 | SEY6210; *HIS3MX6*::$P_{tetoff}$-*GFP-COM2* $P_{YPK1}$*CBSΔ LCB1-13MYC*::*HIS3MX6* | This study |
| RSY2 | SEY6210; *COM2-13MYC*::*HIS3MX6* $P_{YPK1}$*CBSΔ* | This study |
| *com2Δ/com2Δ* | BY4743; *com2Δ*::*KANMX4/com2Δ*::*KANMX4* | Research Genetics |
| *ypk1Δ/ypk1Δ* | BY4743; *ypk1Δ*::*KANMX4/ypk1Δ*::*KANMX4* | Research Genetics |
| *lsp1Δ/lsp1Δ* | BY4743; *lsp1Δ*::*KANMX4/lsp1Δ*::*KANMX4* | Research Genetics |
| *tif3Δ/tif3Δ* | BY4743; *tif3Δ*::*KANMX4/tif3Δ*::*KANMX4* | Research Genetics |
| *stm1Δ/stm1Δ* | BY4743; *stm1Δ*::*KANMX4/stm1Δ*::*KANMX4* | Research Genetics |
| *pcd1Δ/pcd1Δ* | BY4743; *pcd1Δ*::*KANMX4/pcd1Δ*::*KANMX4* | Research Genetics |
| *bmh2Δ/bmh2Δ* | BY4743; *bmh2Δ*::*KANMX4/bmh2Δ*::*KANMX4* | Research Genetics |
| *rim20Δ/rim20Δ* | BY4743; *rim20Δ*::*KANMX4/rim20Δ*::*KANMX4* | Research Genetics |
| *caf20Δ/caf20Δ* | BY4743; *caf20Δ*::*KANMX4/caf20Δ*::*KANMX4* | Research Genetics |
| *yor277cΔ/yor277cΔ* | BY4743; *yor277cΔ*::*KANMX4/yor277cΔ*::*KANMX4* | Research Genetics |
| *rfm1Δ/rfm1Δ* | BY4743; *rfm1Δ*::*KANMX4/rfm1Δ*::*KANMX4* | Research Genetics |
| *tvp18Δ/tvp18Δ* | BY4743; *tvp18Δ*::*KANMX4/tvp18Δ*::*KANMX4* | Research Genetics |
| *sro77Δ/sro77Δ* | BY4743; *sro77Δ*::*KANMX4/sro77Δ*::*KANMX4* | Research Genetics |
| *pin4Δ/pin4Δ* | BY4743; *pin4Δ*::*KANMX4/pin4Δ*::*KANMX4* | Research Genetics |
| *stu2Δ/stu2Δ* | BY4743; *stu2Δ*::*KANMX4/stu2Δ*::*KANMX4* | Research Genetics |

**Table S3. Plasmids used in this study**

| Plasmids | Construct | Reference or source |
|---|---|---|
| pRS415 | *Amp*, *LEU2*, CEN6 | (6) |
| pRS416 | *Amp*, *URA3*, *CEN6* | (6) |
| pRT3 | pRS416; *HIS3MX6*-$P_{ADH1}$-*tTA2*$^{S}$-$P_{tetoff}$-*GFP* | (5) |
| pMT1438 | pDONR221 without BbsI; $P_{SNR52}$-*gRNA*-$T_{SUP4}$ | This study |
| pMT1445 | pRS416; *attR1-Cm*$^{R}$-*ccdB-attR2*-$P_{ADH1}$-$Z_4$EV-$P_{Z4EV}$-*CAS9-CYC1pA* | This study |
| pMT1602 | pMT1438; $P_{YPK1}$-*gRNA* | This study |
| pMT1603 | pMT1445; $P_{YPK1}$-*gRNA* | This study |
| pMT1536 | pRS416; $P_{COM2}$-*COM2*$^{T251A+S370,\ 380A}$-*13MYC* (*COM2* STA3) | This study |
| pMT1547 | pRS416; $P_{COM2}$-*COM2*$^{T251A+S88,\ 164,\ 370,\ 380A}$-*13MYC* (COM2 STA5) | This study |
| pMT1576 | pRS416; $P_{COM2}$-*COM2*$^{T251A+S88,\ 164,\ 235,\ 370,\ 380A}$-*13MYC* (COM2 STA6) | This study |
| pMT1577 | pRS416; $P_{COM2}$-*COM2*$^{T251A+S88,\ 164,\ 235,\ 359,\ 370,\ 380A}$-*13MYC* (COM2 STA7) | This study |
| pMT1578 | pRS416; $P_{COM2}$-*COM2*$^{T251,\ 368A+S88,\ 164,\ 235,350,\ 370,\ 380A}$-*13MYC* (COM2 STA8) | This study |
| pMT1582 | pRS415; $P_{YPK1}$-*YPK1*$^{L424A}$ | This study |
| pYI41 | YEp352 Suppressor #6 (*YPK1*) | This study |
| pYI42 | YEp352 Suppressor #13 (*TIF3*) | This study |
| pYI44 | YEp352 Suppressor #26 (*RIM20* partial, *CAF20*, *HEM14*, *RFM1*) | This study |
| pYI45 | YEp352 Suppressor #28 (*TVP18*) | This study |
| pYI46 | YEp352 Suppressor #34 (*COM2*) | This study |
| pYI47 | YEp352 Suppressor #36 (*SPO77* partial, *PKC1* partial) | This study |
| pYI50 | YEp352 Suppressor #54 (*STM1*) | This study |
| pYI51 | YEp352 Suppressor #9 (*LSP1*) | This study |
| pYI53 | YEp352 Suppressor #17 (*BMH2*) | This study |
| pYI56 | YEp352 Suppressor #56 (*PIN4*) | This study |
| pYI58 | YEp352 Suppressor #6 (*SUT2*) | This study |
| pYI62 | pRS416; $P_{COM2}$-*COM2-13MYC* (*COM2* WT) | This study |
| pYI68 | p$P_{YPK1}$-*LacZ* | This study |
| pRS16 | p$P_{YPK1\text{-}CBS\Delta}$-*LacZ* | This study |
| pRS18 | p$P_{LCB1}$-*LacZ* | This study |
| pRS28 | p$P_{LCB1\text{-}CBS\Delta}$-*LacZ* | This study |
